# Fast and memory-efficient mapping of short bisulfite sequencing reads using a two-letter alphabet

**DOI:** 10.1101/2020.12.21.423849

**Authors:** Guilherme de Sena Brandine, Andrew D. Smith

**Affiliations:** Quantitative and Computational Biology, University of Southern California

## Abstract

DNA cytosine methylation is an important epigenomic mark with a wide range of functions across many organisms. Whole genome bisulfite sequencing (WGBS) is the gold standard to interrogate cytosine methylation genome-wide. Algorithms used to map WGBS reads often encode the four-base DNA alphabet with three letters by reducing two bases to a common letter. This encoding substantially reduces the entropy of nucleotide frequencies in the resulting reference genome. Within the paradigm of read mapping by first filtering possible candidate alignments, reduced entropy of the reference can increase the required computing effort. We introduce another bisulfite mapping algorithm (abismal), based on the idea of encoding a four-letter DNA sequence as only two letters, one for purines and one for pyrimidines. We show that this encoding has greater specificity when subsequences are selected from reads for filtration. Through the two-letter encoding, the abismal software tool maps reads in less time and using less memory than most WGBS read mapping software tools, while attaining similar accuracy. This allows *in silico* methylation analysis to be performed in a wider range of computing machines with limited hardware settings.

## 1 Introduction

DNA cytosine methylation plays a key role in transcription regulation in many eukaryotes, including all mammals. This epigenetic mark is characterized by the addition of a methyl group to the fifth carbon of a cytosine (C), which converts it to a 5-methyl cytosine. With few exceptions, heavily methylated promoters are generally associated with epigenomic repression of the gene. Methylation through distal regulatory regions, such as enhancers, tends to be associated with inactivity of the element. In vertebrates, methylation is added, maintained and removed by enzymatic recognition of cytosines that are directly followed by guanines (Gs). These CpG dinucleotides are the least frequent of all dinucleotides in the genomes of most eukaryotes (often comprising only 1-3% of the sequence). In regulatory regions, however, the frequency of CpGs is significantly higher than the genome-wide average (Saxonov *et al.*, 2006). DNA methylation has been implicated in cell fate decision (Smith and Meissner, 2013), genomic imprinting (Li *et al.*, 1993), X chromosome inactivation (Norris *et al.*, 1991) and retrotransposon silencing (Molaro *et al.*, 2014). Bisulfite sequencing is the gold standard to analyze cytosine methylation (Frommer *et al.*, 1992). Sodium bisulfite treatment converts unmethylated cytosines to uracil while not affecting methylated cytosines. The converted DNA is then amplified in a polymerase chain reaction, which copies uracil as thymine (T). Whole genome bisulfite sequencing (WGBS) couples bisulfite treatment with high-throughput sequencing, which generates hundreds of millions of short reads ranging from 50 to 150 base pairs. WGBS data analysis allows cytosine methylation estimates genome-wide at single-base resolution (Lister and Ecker, 2009; Cokus *et al.*, 2008).

Analysis of short read sequencing data, including WGBS, starts by mapping reads to the reference genome sequence of the organism from which the DNA originates. The problem of read mapping is to find the location in the reference genome that contains the sequence most similar to the read. In general, mapping a read to the reference retrieves the most likely genomic origin for that read. Mappers must be sensitive to differences between read and reference sequences, which may originate from, among other sources, sequencing errors or true genetic variation. The large number of reads necessary to attain sufficient coverage in large genomes requires mapping algorithms to be simultaneously fast and sensitive to various sequence differences between read and reference.

To attain efficient mapping, most short read mapping algorithms start with a filtration step, which reduces comparisons to a small set of reference subsequences that are likely to be optimal candidates. Most filtration methods assume that the read and its optimal mapping location contain a common subsequence. A read is mapped by selecting a set of subsequences from it, then comparing the read to all locations in the reference that match a subsequence. We call subsequences selected from reads for filtration “seeds”. The most similar mapping locations are reported if they attain sufficiently high similarity to the read. The algorithmic efficiency of both filtration and comparison between the read and each candidate globally determine the speed of a mapping algorithm. Reads can originate from either of two complementary genome strands. The most common way to map a read to both strands is to map both the sequence and its reverse-complement to one of the two strands. When WGBS reads are mapped, genome-wide methylation levels can be estimated based on the number of Cs and Ts in reads that map to each C in the reference genome.

Bisulfite conversion makes the problem of mapping WGBS reads different than the traditional DNA read mapping problem. WGBS reads are either T-rich (where most Cs are sequenced as Ts) or A-rich, with the latter occurring when the complementary strand of a converted read is sequenced. We refer to Ts in T-rich reads and As in A-rich reads as bisulfite bases. The reverse-complement of a T-rich read is A-rich, so mapping a T-rich read to the complementary strand is equivalent to mapping an A-rich to the original strand. Multiple four-letter sequences can generate a given T-rich or A-rich sequence once converted, so WGBS mappers must allow both bisulfite bases to each match two bases in the reference to quantify the similarity between a read and a reference sequence.

Several tools exist to map bisulfite-converted reads to a reference genome, with many novel algorithms introduced to address the specific challenges of bisulfite conversion. These tools can be divided in two categories: (i) wrappers of short unconverted DNA read mapping algorithms adapted for bisulfite conversion and (ii) mappers designed specifically for bisulfite-converted reads. Tools within the first category often operate by creating two copies of the reference genome, one that converts all Cs to Ts and one that converts all Gs to As, then mapping the input reads by also replacing any unconverted read Cs with Ts. Bismark (Krueger and Andrews, 2011) and BWA-meth (Pedersen *et al.*, 2014) are two commonly used tools that adopt this approach. Bismark is a wrapper for Bowtie 2 (Langmead and Salzberg, 2012), whereas BWA-meth wraps the BWA-MEM program (Li, 2013). Tools within the second category often incorporate filtration and alignment that account for bisulfite conversion in their implementation, which allows for potential algorithmic optimizations that may result in faster and more memory-efficient mapping times. BSMAP (Xi and Li, 2009) and WALT (Chen *et al.*, 2016) are two commonly used mappers for WGBS data. These two mappers implement similar algorithms, but BSMAP applies filtration by selecting contiguous seeds from the read, whereas WALT uses periodically spaced seeds (Keich *et al.*, 2004). HISAT-3N (Zhang *et al.*, 2021) was developed as a general solution to map reads from sequencing protocols that convert nucleotides. These include, besides WGBS, the SLAM-seq protocol, which converts uracils to Cs to allow the analysis of uracil introduction dynamics in maturing RNAs (Herzog *et al.*, 2017). Bisulfite read mapping algorithms in both categories have varying computational requirements, with more sensitive algorithms usually requiring more time and memory to map reads.

Despite their algorithmic and conceptual differences, a common property of WGBS mappers is filtering candidates by allowing bisulfite bases in seeds to match two possible reference letters, but requiring all other seed bases to match reference letters exactly. Since bisulfite bases account for half of the converted bases, this approach may result in less efficient filtration. In particular, if a seed is selected for filtration based on exact matches, the number of candidates retrieved from the genome will likely increase with the number of bisulfite bases in the selected subsequence, resulting in a greater number of false positive matches. Furthermore, mappers that use the three-letter encoding are typically implemented by keeping two copies of the reference, one for each bisulfite base. In practice, this often requires mappers to use more memory.

Here we show that accounting for the statistical properties of nucleotide frequencies after bisulfite conversion provides an avenue for more efficient filtration and potential improvement in the speed and memory requirement to map WGBS reads. Strand symmetry in eukaryotic genomes results in complementary bases being equally frequent in both strands (Kirkpatrick, 2010). As a consequence of this symmetry, purines (As and Gs) and pyrimidines (Cs and Ts) are also equally represented both before and after bisulfite conversion. This suggests the use of filtration using a two-letter alphabet encoding, which simultaneously converts purines to one letter and pyrimidines to another letter. Encoding both reads and reference in this alphabet allows filtration in both T-rich and A-rich reads. We show that, compared to the commonly adopted three-letter encoding, filtration using the two-letter alphabet increases specificity when a fixed number of bits is available to encode an arbitrary subsequence selected from a read. We describe theoretical results for fingerprint-based filtration (Karp and Rabin, 1987) in genomes formed by independent and identically distributed (*i.i.d*) letters and show that many eukaryotic genomes have comparable properties to those derived theoretically assuming *i.i.d* sequences. We introduce an implementation of this approach as a novel algorithm and software tool, named abismal, that is optimized for WGBS read mapping. Using publicly available data, we show that abismal attains equivalent accuracy to commonly used WGBS mappers in scientific research, but requires significantly less time and memory to map reads while attaining comparable results.

## 2 Methods

### 2.1 Indexing with a two-letter alphabet

Mapping bisulfite sequencing reads typically involves converting the genome into a three-letter sequence, with C→T or G→A to simulate the bisulfite conversion process. Our approach is motivated by the observation that if the reads and the genome are converted into two-letter sequences, with both C→T and G→A simultaneously, a match in the two-letter converted sequences is a necessary condition for there to be a match between the three letter sequences. Moreover, the same two-letter encoding works for both T-rich reads and A-rich reads. If this necessary condition is specific enough, and if it can be tested efficiently, the approach may have advantages. Beyond the ability to use the same encoding to map both A-rich and T-rich reads, we present statistical rationale in the next section for the benefits of this strategy.

We define a *k*-mer as a sequence of *k* consecutive letters. We encode sequences by converting each *k*-mer to a numerical representation that uses one bit per position, distinguishing purines from pyrimidines. In particular, with *h*_2_(A) = *h*_2_(G) = 0 and *h*_2_(C) = *h*_2_(T) = 1, the two-letter fingerprint for a *k*-mer *w* is defined as

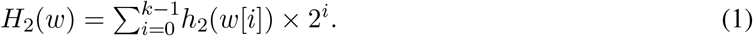

Under this scheme, *k*-mers over the original DNA alphabet are associated with *k*-bit binary numbers, and 2^*k*^ distinct DNA sequences have the same fingerprint. This same strategy can be applied in the context of spaced seeds (Ma *et al.*, 2002), encoding ordered but non-contiguous sets of *k* letters. The advantages of spaced seeds are most pronounced when they span large portions of the read, which renders them less sensitive to insertions and deletions (indels). We restrict our analyses to contiguous sequences.

### 2.2 Theoretical analysis of the two-letter encoding

Here we analyze the theoretical efficiency of the two-letter fingerprint strategy, defined in equation (1), as a function of the number of bits in the fingerprint. This will be done by comparison with the three-letter encoding used in most bisulfite sequencing mappers, which reflects the chemical process of bisulfite conversion. In particular, the three-letter strategy encodes letters as *h*_3_(A) = 0, *h*_3_(G) = 1 and *h*_3_(C) = *h*_3_(T) = 2. This encoding simply equates T and C. Using *h*_3_, the fingerprint for sequence *w* can be constructed as

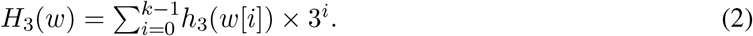

The function *H*_3_ evaluates a radix-3 polynomial, while in practice it has been more common to use a radix-4 variation, defined as

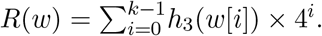

The two main distinctions between radix-3 and radix-4 are (i) the size, in bits, for the resulting fingerprint value, and (ii) the operations required to evaluate the fingerprint function. Note that *R* can be evaluated using logical operations, while *H*_3_ requires multiplication and modular arithmetic. Since *H*_3_(*w*) = *H*_3_(*w*′) if and only if *R*(*w*) = *R*(*w*′), the radix-3 and radix-4 encodings are equivalent with respect to accuracy characteristics as fingerprint functions, but when bounded by a common maximum value, the image of the radix-3 encoding contains more elements.

As outlined in the previous section, the time required by a mapping algorithm depends on the total number of fingerprint hits, which, for a particular fingerprinting scheme, is proportional to the expected hit rate for a random *k*-mer. We assume a random genome sequence of infinite size with *i.i.d* letters, and will address relaxations of this assumption later. We further assume that Pr(A) = Pr(T) = *p* and Pr(C) = Pr(G) = *q*, reflecting strand symmetry in eukaryotic genomes. Importantly, this symmetry ensures that *p* + *q* = 1/2. We define the expected hit rate of a fingerprint function as the expected fraction of positions whose fingerprint is *H*(*w*), where *w* is a uniformly sampled *k*-mer from the genome. This emulates the process of selecting fingerprints based on seeds from a large number of reads that were themselves sampled from the reference sequence, as is the case with real datasets (with the exception of sample contamination). The expected hit rate *Z* can be calculated as

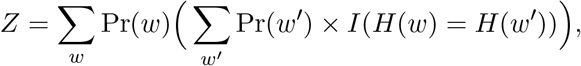

for indicator function *I*. If *k*-mers *w* and *w* are sampled at random from the genome, this is equivalent to

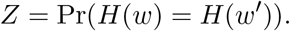

If we assign fingerprints to each genome position using two functions *H* and *H* with the same codomain Ω, the one with lower expected hit rate is preferred. For a fixed reference genome, let *u* be a fingerprint and *n*(*u*) be the number of *k*-mers whose fingerprint is *u*. The empirical expected hit rate can be calculated by

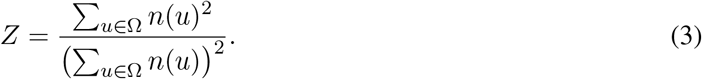

We will also define the theoretical hit rate *z* to be the expected hit rate in the specific case of infinite *i.i.d.* genomes where *p* + *q* = 1/2. We can directly devise bounds for the theoretical hit rate under the two-letter encoding *z*_2_ and three-letter encoding *z*_3_.

First, consider the two-letter fingerprint strategy *H*_2_ defined in equation (1). Our statistical assumptions on the base frequencies in the genome imply that the theoretical hit rate is *z*_2_ = (1/2)^*k*^ regardless of the values of *p* and *q*. We can also deduce that the lowest possible theoretical hit rate *z*_3_ for *H*_3_ is (3/8)^*k*^. This can be proven as follows. Consider *w* and *w* to be two independently sampled *k*-mers. The independence allows the following rearrangement

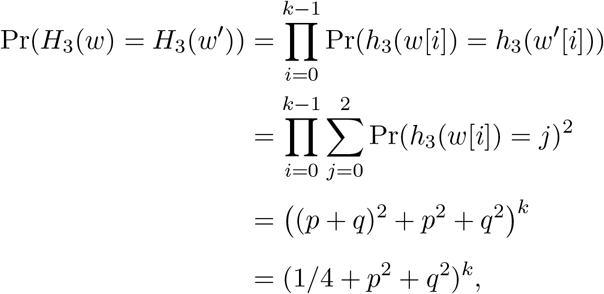

by our assumption that *p* + *q* = 1/2. The minimal value of (3/8)^*k*^ is attained when *p* = *q* = 1/4.

If we evaluate fingerprints for *k*-mers within reads, the specificity is greater for the three-letter strategy, which should be intuitive. However, our capacity to index fingerprint values for efficient retrieval depends on the number of distinct values taken by the fingerprint function. In other words, it is more appropriate to compare the two strategies when the number of fingerprint values is fixed, rather than the number of letters in the fingerprint.

Assume fingerprint values are *b*-bit non-negative integers, so fingerprints are values between 0 and 2^*b*^ − 1. For the two-letter encoding, *b* letters can be encoded using *b* bits. For the three-letter encoding, the number of letters that can be represented using *b* bits is ⌊*b/* log_2_(3)⌋. In this case, the expected hit rate is lower for the two-letter encoding. This can be proven by analyzing the ratio between *z*_2_ and the lowest possible value of *z*_3_, since

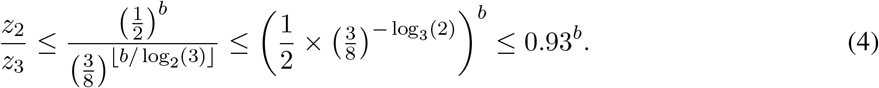

Equation (4) indicates that any specificity lost in the two-letter encoding, compared with the three-letter encoding, may be regained by increased uniformity in the distribution of fingerprint values (Figure 1A). This result, however, assumes *k*-mers are sampled with *i.i.d* letters. Consider a genome containing an extreme abundance of poly-purine sequences. All *k*-mers sampled from this genome would have the same fingerprint under *H*_2_ but would be distinguishable under *H*_3_. We can quantify, using real genome sequences, how their empirical expected hit rates deviate from *i.i.d.* results using equation (3). We compared the empirical and theoretical hit rates for six species: *H. sapiens*, *M. musculus*, *D. rerio*, *G. gallus*, *P. troglodytes* and *A. thaliana*, both under the two-letter and three-letter encodings. For 2 ≤ *b* ≤ 28, and in all six genomes, the empirical and theoretical ratios between expected hit rates are below the bound derived in equation (4) (Figure 1B), which suggests that the *k*-mer distribution in the genomes analyzed favors specificity in the two-letter encoding at least as much as in genomes generated by large *i.i.d* sequences. This means that the two-letter encoding can be used as an efficient filtration strategy for read mapping in these genomes.

**Figure 1:**
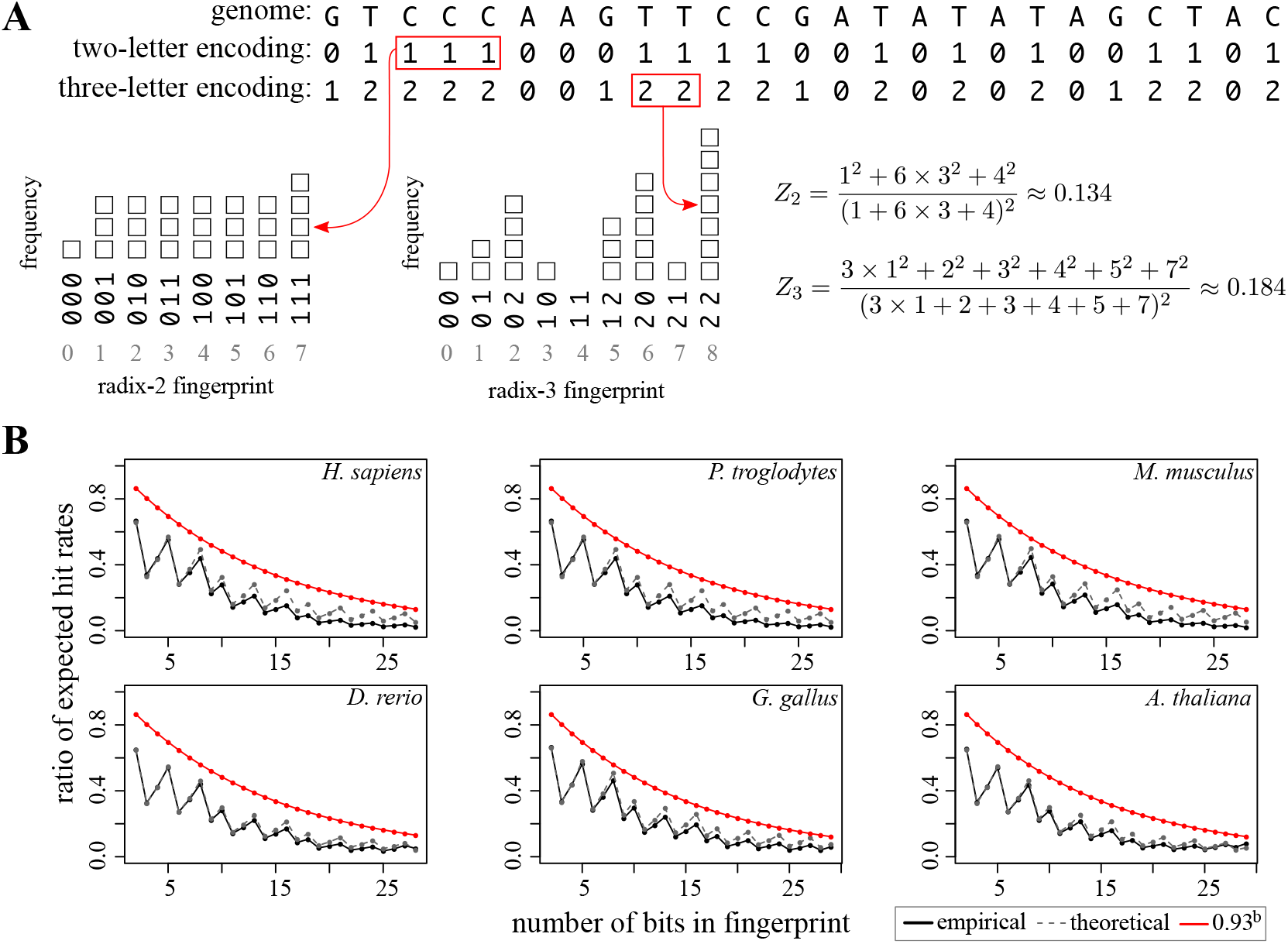
Theoretical and empirical comparison between two-letter and three-letter encodings. **(A)** An example comparison between the expected hit rates. A small *i.i.d.* genome of size 25 is shown, as well as its two-letter and three-letter conversions, according to equations (1) and (2). Fingerprints can take values in Ω = {0, · · · , 8}, and each value is represented as *k*-mers both in radix-2 and radix-3 encodings. The expected hit rates for both encodings *Z*_2_ and *Z*_3_ are calculated using equation (3). Red boxes show example *k*-mers that are counted in both encodings. **(B)** Comparison between the ratio of empirical expected hit rates *Z*_2_*/Z*_3_ and the theoretical ratio *z*_2_*/z*_3_, which assumes infinite *i.i.d.* genomes. Empirical expected hit rates *Z*_2_ and *Z*_3_ were calculated using equation (3). The *k*-mer sizes are given by *k*_2_ = *b* and *k*_3_ = ⌊*lb/* log2(3)⌋, where *b* is the number of available bits, ranging from 2 to 28.

### 2.3 Implementation choices

The two-letter encoding, like its three-letter counterpart, can be incorporated as an intermediate step in any DNA read mapping algorithm adapted for bisulfite sequencing reads. Wrappers for three-letter encoding adaptations of existing DNA read mapping software often operate by creating two indices of the original genome that simulate bisulfite conversion, then mapping reads using the resulting indices and combining results. It is more challenging, however, to directly write wrappers around existing DNA read mapping code to use the two-letter encoding. This is because encoded reads are used for filtration, but both read and reference must be available as four-letter sequences in order to calculate alignment scores. To demonstrate the usage of the two-letter encoding for WGBS read mapping, we introduce another bisulfite mapping algorithm (abismal), a specific implementation of a bisulfite sequencing mapper that uses the two-letter encoding as a filtration strategy. For read *r* = *r*_1_ … *r*_*n*_, we select *k*-mers from *r* and retrieve all positions in the genome where the two-letter encoding of each *k*-mer occurs. Each of these positions is assessed by scoring the alignment of *r* with the corresponding genomic sequence. The position having the best alignment score is retained as the mapping location for *r*, and if two locations share the best score, the read is considered to map ambiguously. Most algorithms for mapping short reads fall within this framework. One of the most significant factors in the time required by this framework is the number of full comparisons that must be done for each read. We outline several important implementation choices below.

Our implementation uses a direct address table to associate fingerprints with positions in the genome. Before mapping reads, the genome is indexed through a counting sort algorithm (Seward, 1954; Knuth, 1997). The indexing algorithm sorts a subset of genome positions in a direct address table based on the two-letter *k*-mers starting at each position. Initially, the frequency of all 2^*k*^ two-letter *k*-mers is counted in a linear pass through the selected genome positions and stored in a count table, which is then transformed into a cumulative sum vector of size 2^*k*^ + 1. A second pass through the genome populates the direct address table using the cumulative sum vector to retrieve the proper index location of each genome position. Reads are mapped by selecting a set of *k*-mers from it, then aligning the read to every indexed genome position that match the selected *k*-mers exactly. The resulting count table from the indexing step allow positions to be retrieved based on fingerprint values. The four-letter read is aligned to positions in the four-letter reference, with bisulfite bases in reads matching two possible reference bases. The best alignment is identified by locating the best Hamming distance matches (not allowing indels) and then computing a banded Smith-Waterman score for each of them (Smith and Waterman, 1981). The full description of the abismal algorithm, as well as a more detailed rationale on the choice of parameters used and their effects in mapping accuracy is provided in the Supplementary Methods.

## 3 Results

The mapped reads are used to estimate the methylation level at each CpG site (or possibly each cytosine) in the genome. The methylation level at a site is the fraction of original molecules (*e.g.* 2 per cell for a diploid sample) carrying the methyl mark at that site. This situation is similar to genotyping a site using mapped reads from a whole genome sequencing experiment. Unlike genotype, methylation levels may take values beyond the 1.0 and 0.5 frequencies associated with homozygous and heterozygous sites, and may instead take any value between 0 and 1. Consequently, systematic errors in mapping may lead to incorrect biological conclusions.

### 3.1 Comparison criteria

To evaluate performance, we used both simulated data and 15 public high-quality datasets obtained from the Sequence Read Archive (Leinonen *et al.*, 2010). These datasets include single-end and paired-end reads from traditional WGBS protocol (*i.e.* with T-rich reads on the first end and A-rich reads on the second end) and from the random-priming post-adapter bisulfite tagging (RPBAT) protocol (Miura and Ito, 2015). RPBAT is an amplification-free protocol that randomly sequences T-rich and A-rich reads with the property that paired ends have complementary bisulfite bases. Traditional single-end (Brocks *et al.*, 2017; Manakov *et al.*, 2015; de Mendoza *et al.*, 2020; Zhang *et al.*, 2017; Shahryary *et al.*, 2020) and paired-end (Do *et al.*, 2020; Decato *et al.*, 2017; Kamstra *et al.*, 2018; Lee *et al.*, 2017; Yong-Villalobos *et al.*, 2015) datasets were selected from human and four model organisms (mouse, chicken, zebrafish and arabidopsis), whereas RPBAT datasets (Miura and Ito, 2015; Guo *et al.*, 2015; Miura *et al.*, 2019; Leng *et al.*, 2019; Bian *et al.*, 2018) originate from human samples (Supplementary Table S1). We show performance results on three distinct metrics: simulation accuracy (in the form of sensitivity and specificity), number of reads (or read pairs) mapped at least once, and average error rate between read and reference.

Simulation accuracy for reads sampled with error from the reference genome is an objective assessment of mapper performance. Mapped reads can be compared to their true locations of origin to assess the sensitivity and specificity of each mapper. Sensitivity is defined as the fraction of correctly mapped reads relative to the total number of simulated reads, and specificity is the fraction of reported reads that are correctly mapped. To quantify both metrics, paired-end reads were simulated, both in traditional and RPBAT modes, using Sherman (Krueger, 2017) and setting various error rates from 0% to 5%. We also simulated 1% and 70% methylation levels for cytosines outside and inside of the CpG context, respectively. These methylation parameters simulate well-established methylation rates from most healthy mammalian somatic cells estimated from WGBS data with high conversion rate (Bernstein *et al.*, 2010). We varied read lengths from 50 to 150, a range that encompasses most read lengths generated by Illumina sequencers.

For real data, the true location of origin for each read is not known, but metrics can be used to assess the relative quality of mapper outputs. The percentage of reads mapped at least once is an estimate of mapper sensitivity. In particular, paired-end reads that map concordantly (here defined as at most 3000 bases apart and in opposite strands), are likely to be correct candidates. Many mappers, including abismal, map each end of read pairs independently. Since all tested reference genomes are large (the smallest with 135 Mbp), it is unlikely that reads incorrectly mapped are concordant by chance alone. To estimate specificity, we measured the error rate of mapped reads, defined as the ratio between the total number of edits (mismatches, insertions and deletions) and the total number of bases aligned to the reference. We expect more specific mappers to have lower error rates.

We interpreted abismal’s accuracy, time and memory metrics by comparing it with five WGBS mappers. In addition to abismal, we mapped simulated and real reads with Bowtie 2 (through Bismark), BSMAP, BWA-MEM (through BWA-meth), HISAT-3N and WALT. We mapped RPBAT datasets using abismal, Bismark and BSMAP, since these are the only mappers with modes that allow T-rich and A-rich reads to be mapped interchangeably. Parameters for mappers were chosen to maximize the similarity between algorithms (*e.g.* similar alignment scores and similarity cut-offs to consider a read “mapped”). We ran each mapping tool on machines with identical hardware, system software and user environments. We instructed mappers to use 16 threads, which favors mappers that make efficient use of parallelism. Parameters for each mapper and hardware settings used in every test are detailed in the Supplementary Methods.

### 3.2 Comparison results

In simulated reads, HISAT-3N attains highest sensitivity and BWA-meth attains highest specificity, with abismal closely approximating each mapper in both criteria. Abismal’s overall sensitivity and specificity are lower for 50 and 60 bp reads, but steadily increase from 70 to 150 bp. The lower accuracy for shorter reads stems from the fact that two-letter seeds in abismal are larger (24 bp) compared to other mappers (*e.g.* 22 bp for Bismark and 16 bp for BSMAP), so they often cover a high fraction of the read and mismatches in certain locations may compromise all sampled seeds. Conversely, in higher error settings, two-letter seeds may be invariant to certain mutations, and seeds with errors may still retrieve the correct candidate. The simulations results are consistent in the traditional paired-end setting (Supplementary Figure S1) and RPBAT setting (Supplementary Figure S2). In real data, abismal, Bismark, BSMAP and HISAT-3N have the most mapped reads (Figure 2B; Supplementary Table S2), whereas abismal, WALT and BWA-meth show the lowest error rates (Figure 2C; Supplementary Table S2). WALT attains higher specificity due to its spaced seed strategy, which disallows mismatches in the most error-prone 3’ end of short reads. BWA-meth’s higher specificity stems from its maximum exact matching paradigm, which retains candidates based on the largest contiguous subsequence that matches some region of the genome exactly. Overall, abismal attains similar metrics to the best-performing mapper in all three comparison criteria.

**Figure 2:**
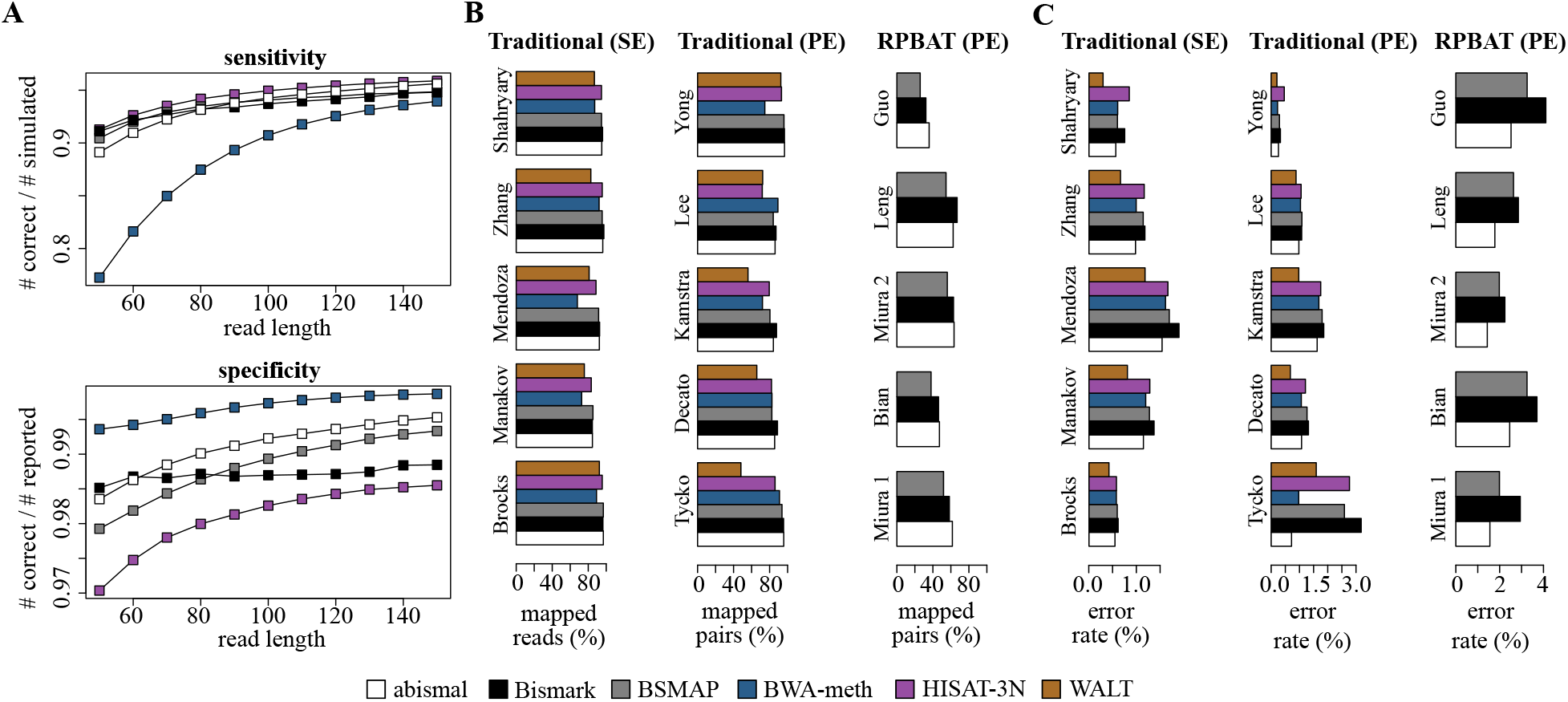
Accuracy comparison of mapper outputs. **(A)** comparison of mapping sensitivity (top) and specificity (bottom) for simulated reads with 0 to 5% average error rate and bisulfite conversion of 99% for non-CpGs and 30% for CpGs. Sensitivity is measured by the fraction of simulated reads correctly mapped, whereas specificity is defined by the fraction of reported reads that are correctly mapped. **(B)** estimated sensitivity for real data, defined by number of reads (for SE datasets) or concordant pairs (for PE datasets) mapped at least once. **(C)** estimated specificity for real data as the average edit distance of mapped reads, defined by the ratio between mismatches + indels and the total number of read bases mapped to the reference.

We further compared mapper outputs through several methylation-specific summary statistics on read bases that map to cytosines, and specifically CpGs, in the genome. We quantified average methylation levels in three different ways: mean methylation, weighted mean methylation and fractional methylation, as previously suggested (Schultz *et al.*, 2012). We also measured the fraction of covered bases across the genome and the bisulfite conversion rate, estimated by the fraction of unconverted cytosines mapped to Cs outside the CpG context. The results show that these summary statistics, in both cytosines and CpGs, are nearly identical across the four mappers, with no clear biases in any method (Supplementary Table S2).

The mapping time comparison shows that abismal is generally faster, with HISAT-3N and BWA-meth attaining similar speed in some human samples (Figure 3A; Supplementary Table S3). For RPBAT datasets, abismal attains both higher mapping speeds and sensitivity than Bismark and BSMAP. Finally, abismal’s memory requirement to map reads is substantially lower than other mappers (Figure 3B; Supplementary Table S3). For the human genome, abismal requires about 3.5 GB of RAM to map reads and 4.5 GB to index the genome, whereas HISAT-3N, the second most memory-efficient mapper, requires 8 GB. Bismark and BWA-meth use more memory because they operate by running several parallel instances of Bowtie 2 and BWA-MEM, respectively, for the various bisulfite conversion combinations of the reference genome. This means that the memory required to map reads scales with the number of threads used. The low memory use attained by abismal stems from two implementation choices made possible by the two-letter encoding (i) the property that only one strand of the reference needs to be indexed and stored in memory and (ii) the higher uniformity of two-letter *k*-mers relative to the three-letter alphabet (Supplementary Figure S3), which allows efficient usage of minimizers to index positions and select *k*-mers from reads (Roberts *et al.*, 2004). Indexing only minimizers significantly decreases the size of the direct address table (Supplementary Methods).

**Figure 3:**
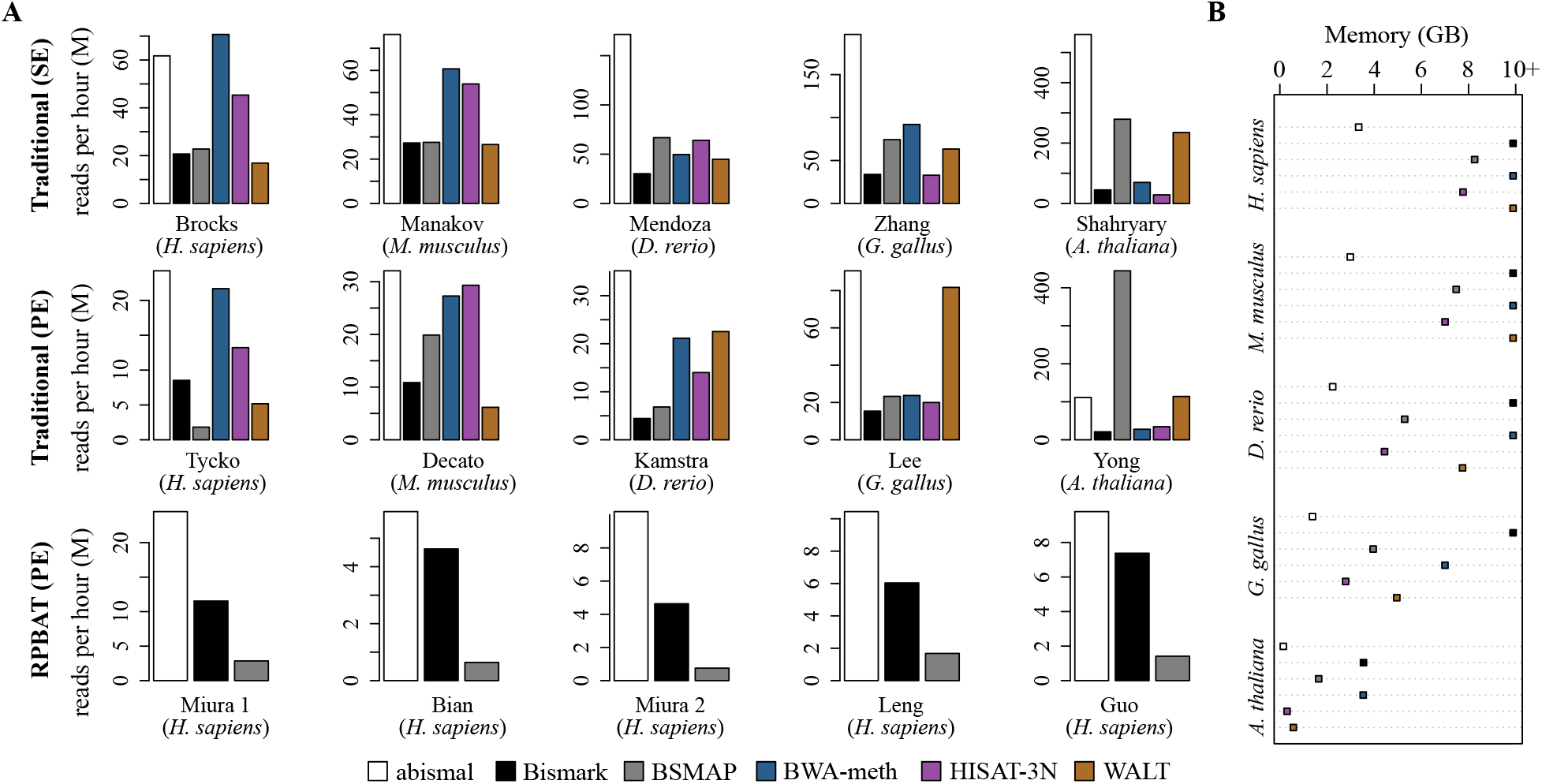
Speed and memory usage comparison between mappers for traditional single-end (SE), paired-end (PE) and RPBAT samples. **(A)** Number of reads (in millions) mapped per hour for each dataset selected for comparison. For abismal and BSMAP, the total mapping time considered is given by the sum of indexing and mapping times. **(B)** Maximum memory used to map each paired-end sample. Memory requirements higher than 10 GB are capped at the 10+ GB mark in the x axis.

## 4 Discussion

We demonstrated the benefits of using a two-letter encoding to map bisulfite sequencing reads and its effect on mapping efficiency. The equal frequency of purines and pyrimidines in eukaryote genomes makes this encoding useful for filtration in bisulfite sequencing read mapping algorithms. The two-letter encoding can be used in any filtration method that selects subsequences from a read as exact matches to the reference.

The abismal software tool maps reads with comparable sensitivity to several commonly used mappers in less time and uses significantly less memory. Since most downstream methylation analyses, such as individual cytosine methylation estimates, require less time and memory than mapping reads (Song *et al.*, 2013), abismal makes basic *in silico* methylation data analysis feasible in machines with low computing power.

There are many avenues for speed and memory use improvements in the abismal algorithm. Abismal uses a random ordering of two-letter *k*-mers to select minimizers in both reads and reference (Marçais *et al.*, 2017). Accounting for the frequency of *k*-mers within a genome may provide opportunities to index less frequent *k*-mers and decrease the number of false-positive matches (Jain *et al.*, 2020). Other genome-aware orderings may also lead to fewer indexed positions and thus less memory use in larger genomes (Orenstein *et al.*, 2017). Finally, improvements in computer architecture continuously lead to increased efficiency in pre-alignment filtration (Alser *et al.*, 2020) and speed of Smith-Waterman local alignments (Zhao *et al.*, 2013), both of which are important steps in most short read mapping algorithms, including abismal.

## 5 Data availability

Abismal is available on GitHub (https://github.com/smithlabcode/abismal). It is a free Open Source software tool under the GNU Public License (GPL) version 3.0. SRA accessions of tested datasets are available in Table 1.

**Table 1:** Datasets used for accuracy, speed and memory comparisons between mappers.

| Dataset | Accession | Species | Phenotype | Protocol | Reads / Pairs | Read length |
| --- | --- | --- | --- | --- | --- | --- |
| Brocks (Brocks *et al.*, 2017) | SRR3498383 | *H. sapiens* | H1299 cell line | Traditional (SE) | 163,725,478 | 100 |
| Manakov (Manakov *et al.*, 2015) | SRR2096734 | *M. musculus* | Spermatocytes | Traditional (SE) | 308,662,065 | 100 |
| Mendoza (de Mendoza *et al.*, 2020) | SRR10606701 | *D. rerio* | Forebrain | Traditional (SE) | 168,367,763 | 96 |
| Zhang (Zhang *et al.*, 2017) | SRR5015166 | *G. gallus* | Breast muscle | Traditional (SE) | 139,994,176 | 125 |
| Shahryary (Shahryary *et al.*, 2020) | SRR12075121 | *A. thaliana* | Leaf | Traditional (SE) | 76,256,688 | 151 |
| Tycko (Do *et al.*, 2020) | SRR10165530 | *H. sapiens* | Liver | Traditional (PE) | 52,607,981 | 144 |
| Decato (Decato *et al.*, 2017) | SRR3897178 | *M. musculus* | Whole placenta | Traditional (PE) | 124,995,278 | 90 |
| Kamstra (Kamstra *et al.*, 2018) | SRR5756263 | *D. rerio* | Whole embryo | Traditional (PE) | 121,650,843 | 150 |
| Lee (Lee *et al.*, 2017) | SRR5644125 | *G. gallus* | Retina | Traditional (PE) | 59,768,935 | 125 |
| Yong (Yong-Villalobos *et al.*, 2015) | SRR2296821 | *A. thaliana* | Pit root | Traditional (PE) | 21,543,631 | 90 |
| Miura 1 (Miura and Ito, 2015) | DRR019425 | *H. sapiens* | K562 cell line | RPBAT (PE) | 69,919,315 | 100 |
| Bian (Bian *et al.*, 2018) | SRR7461526 | *H. sapiens* | CCL23 cell line | RPBAT (PE) | 19,986,722 | 150 |
| Miura 2 (Miura *et al.*, 2019) | SRR7757863 | *H. sapiens* | IMR-90 cell line | RPBAT (PE) | 99,575,506 | 151 |
| Leng (Leng *et al.*, 2019) | SRR9646129 | *H. sapiens* | Single-cell 8-cell PGC | RPBAT (PE) | 71,902,273 | 126 |
| Guo (Guo *et al.*, 2015) | SRR2013804 | *H. sapiens* | Single-cell 10-week PGC | RPBAT (PE) | 91,262,329 | 125 |

