## Supplementary figures and methods for "Fast and memory-efficient mapping of short bisulfite sequencing reads using a two-letter alphabet"

Guilherme de Sena Brandine          Andrew D Smith

#### Supplementary Figures

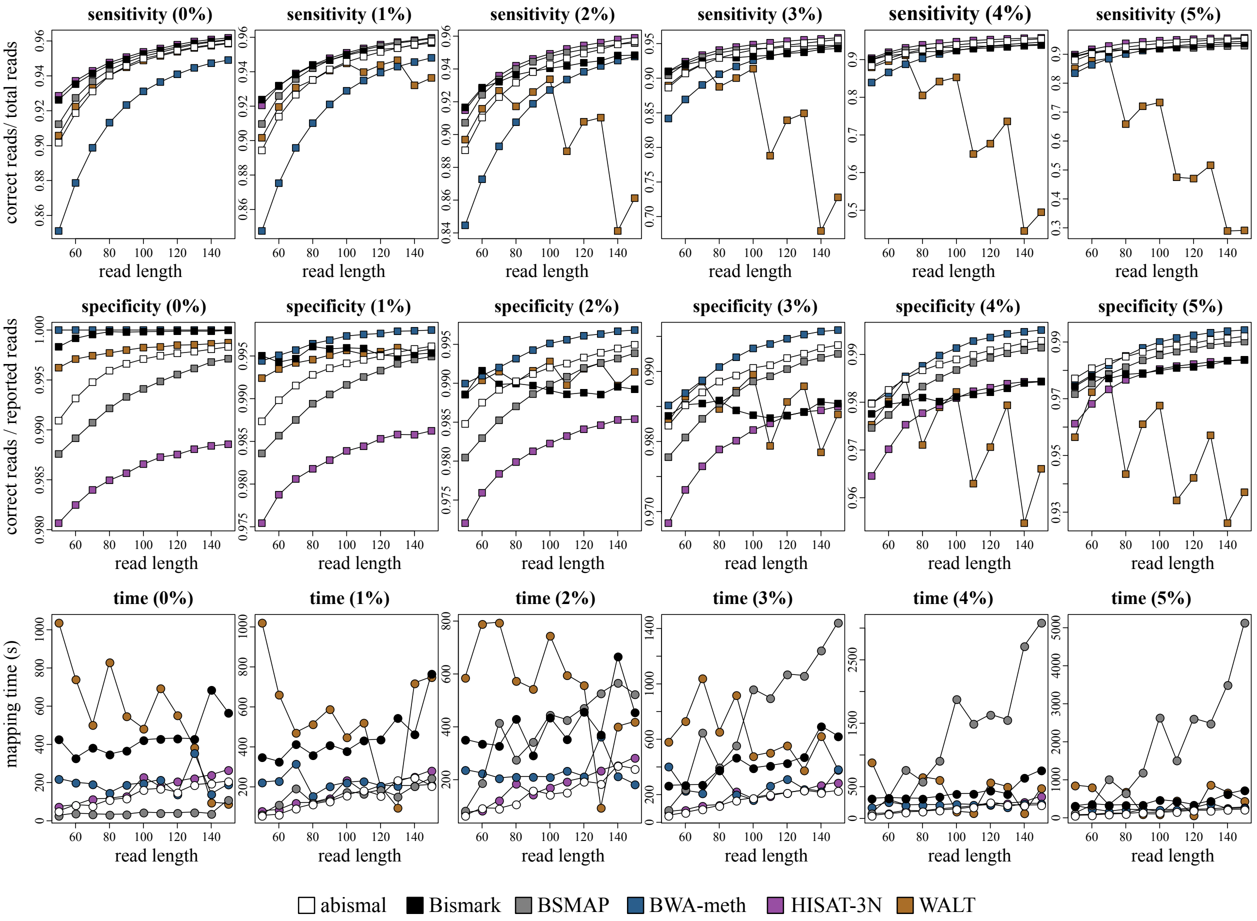


Supplementary Figure S1: Sensitivity, specificity and mapping time for 1 million simulated paired-end reads from the human genome under various read lengths. Error rates are defined in parentheses, varying from 0 to 5%.

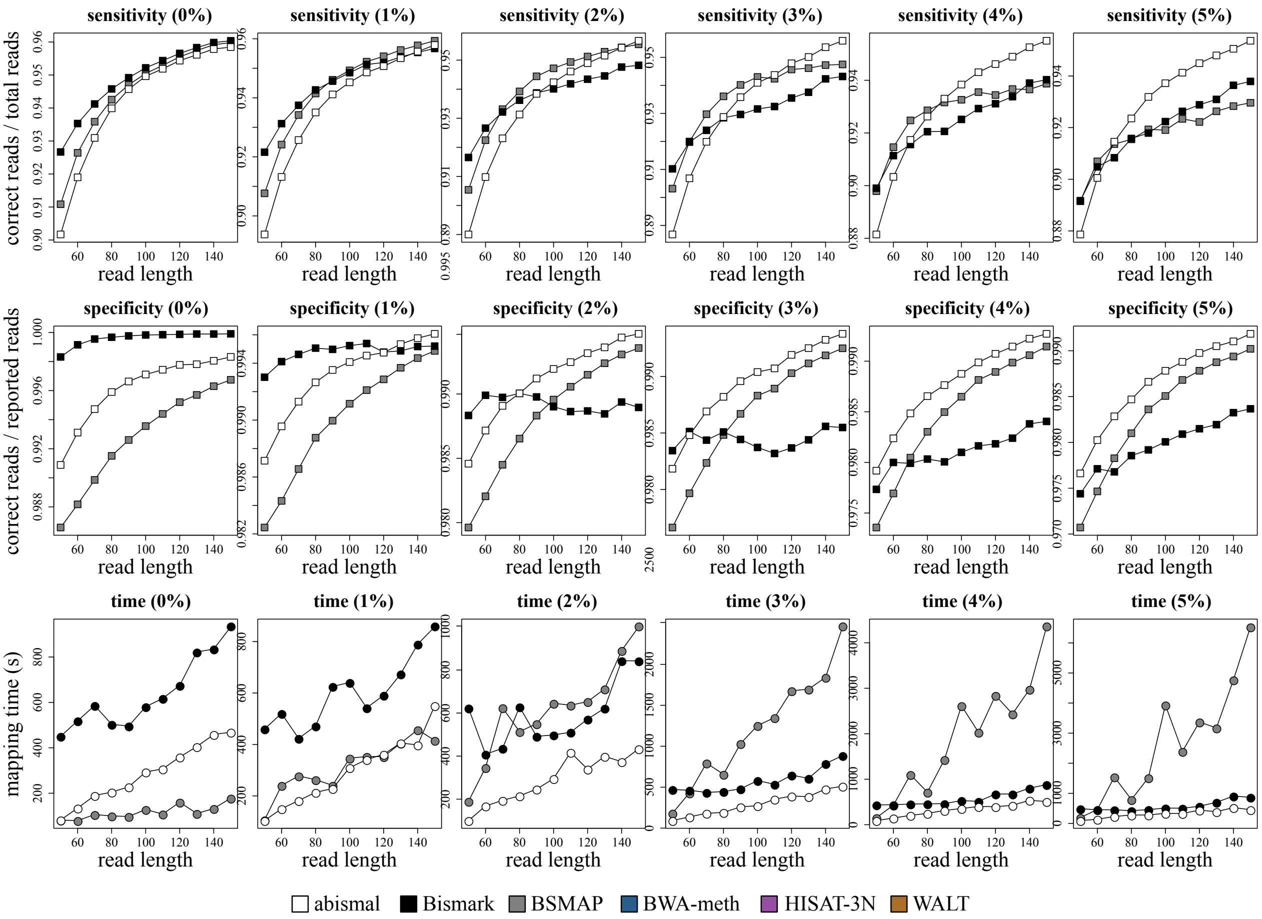


Supplementary Figure S2: Sensitivity, specificity and mapping time for 1 million simulated paired-end RPBAT reads from the human genome under various read lengths. Error rates are defined in parentheses, varying from 0 to 5%.

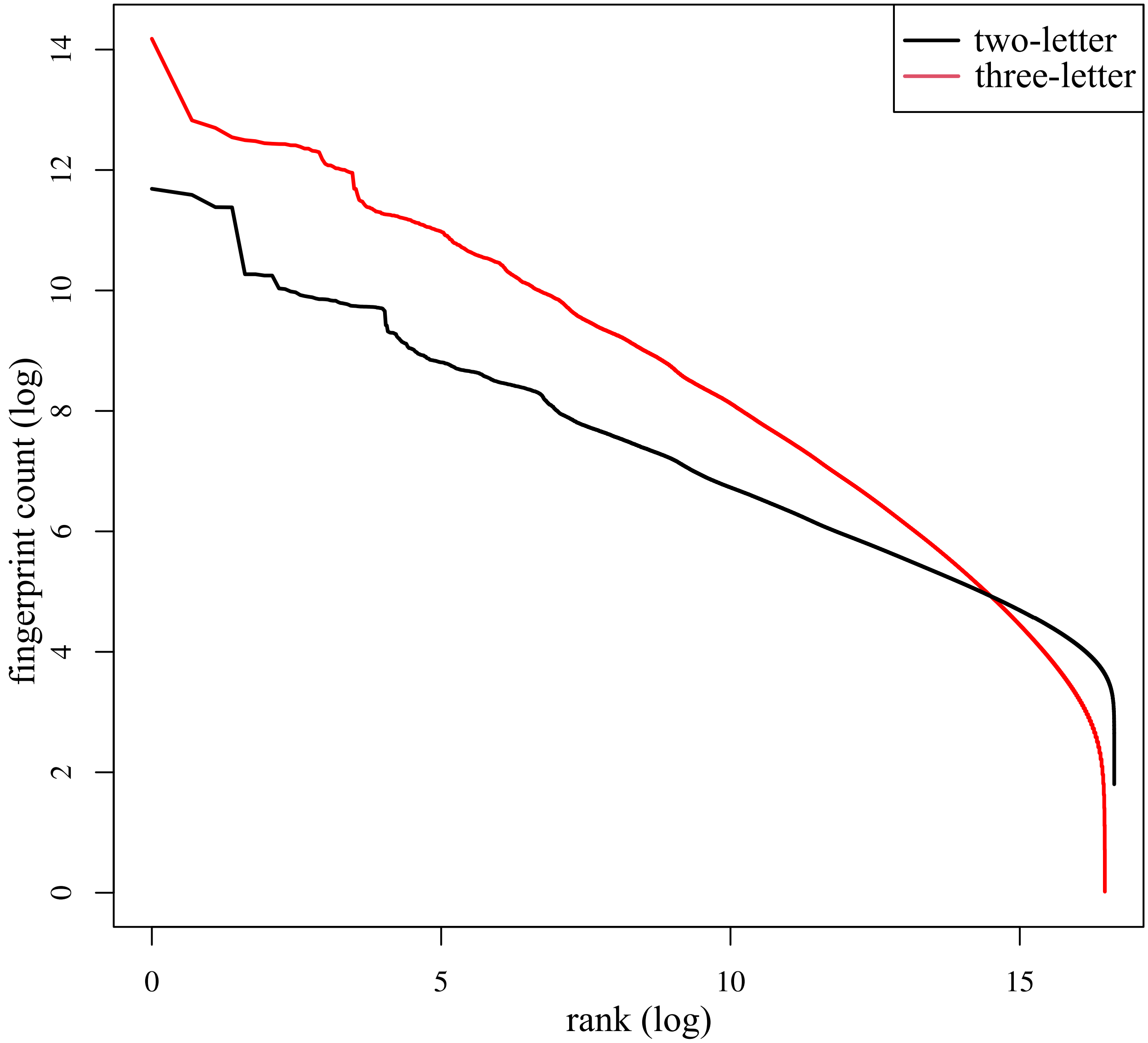


Supplementary Figure S3: Comparison of fingerprint frequencies for the two-letter and three-letter encodings in the human genome. Fingerprints were counted using $b = 24$ bits, so values range from $0$ to $2^b - 1$. Fingerprints of every contiguous sequence under the two- and three-letter alphabets were counted and sorted in decreasing order. The $k$-mer sizes are $k_2 = 24$ for the two-letter encoding and $k_3 = \lfloor 24/\log_2(3) \rfloor = 15$ for the three-letter encoding.

### Supplementary Methods

#### 1 The abismal algorithm

##### 1.1 Reference genome indexing

Indexing of an input genome sequence of length $M$ is performed by grouping positions based on the two-letter $k$-mer sequence starting at each position. We will define the $k$-mer sequence of a position $i$, with $0 \leq i \leq M - k$ to be the two-letter sequence of the $k$-mer starting at position $i$ and ending at position $i + k - 1$. A fixed value of $k$ is assumed given, and it is set to $k = 128$ in the abismal software tool.

We seek to build a genome index, defined as an ordered subset of $\{0, \ldots, M - k\}$, with positions grouped together based on identical $k$-mer sequences. The abismal index is built in four steps: (1) minimizer selection, (2) determining fingerprint frequencies, (3) constructing the initial index and (4) final sorting. The first step partitions genome positions according to $k$-mers that are minimizers for some window of fixed length containing that position. The second and third steps group genome positions by placing locations with identical $k$-mers together. This is done through a counting sort algorithm for fingerprints of smaller $l$-mers, where $l \leq k$. The fourth step further sorts positions based on the subsequent $k - l$ bases. When mapping reads, filtration based on a given two-letter sequence of length $k'$, for $l \leq k' \leq k$, is performed by direct look-up on the first $l$ bases and subsequent binary search of each of the $k' - l$ bases.

**First step: minimizer selection** In the first step, a boolean array $b$ of size $M - k + 1$ is created, where $b_i = 1$ if position $i$ will be indexed in the genome and $b_i = 0$ otherwise. We fill $b_i$ by selecting minimizers in every window of size $W$. We first define a random ordering of all the $2^l$ possible $l$-mers. Then, for $0 \leq j < M - k + 1 - W$, we select position $i \in \{j, \cdots, j + W - 1\}$ if the $l$-mer starting at position $i$ is the minimum of all $l$-mers starting at any position in the interval. We set $b_i = 1$ if and only if $i$ is the minimizer of some window. This procedures guarantees that at least one base in every contiguous window of length $W$ has $b_i = 1$. This also means that the distance between two consecutive positions where $b_i = 1$ is at most $W$. In the subsequent steps, we will only consider the subset of positions in the genome where $b_i = 1$. We will say position $i$ is indexed if $b_i = 1$ and non-indexed otherwise.

**Second step: determining fingerprint frequencies** In the second step, a vector of size $2^l + 1$ is built storing the counts of all $2^l$ contiguous two-letter $l$-mers across the selected minimizer positions. Formally, for $0 \leq v \leq 2^l - 1$, we define $\kappa(v)$ to be the number of times a two-letter sequence with binary representation $v$ has been seen in any position $i$ where $b_i = 1$. Denoting $\gamma' = (\kappa(0), \ldots, \kappa(2^l))$, we create the cumulative sum vector of $\gamma$ of $\gamma'$, defined by $\gamma_i = \sum_{j=0}^{i-1} \gamma'_j$, which will be used in the second step to create the sorted vector based on the first $l$ letters, and also when mapping reads. The vector $\gamma$ is such that $\gamma_0 = 0$ and $\gamma_{2^l}$ is the total number of indexed bases in the first step.

**Third step: index construction** In the third step, a direct address table $\alpha$ of unsigned 32-bit positions of size $\gamma_{2^l}$ is allocated, and each genome position with $l$-mer sequence $v$ is placed in the table in position $\gamma_v$, that is, if position $i$ has $l$-mer $v$, then $\alpha(\gamma_v) = i$. After position $i$ is added to $\alpha(\gamma_v)$ , the value of $\gamma_v$ is decremented, so the next time a position $j$ with $l$-mer $v$ is seen, it will be placed in $\alpha$ preceding $i$. After decrementing all occurrences of each $l$-mer $v$, the list of positions with two-letter $l$-mer $v$ in the genome are between elements $\alpha(\gamma_v)$ and $\alpha(\gamma_{v+1} - 1)$. This allows the resulting table $\gamma$ to be used when mapping reads to retrieve all positions in the genome with a given $l$-mer sequence using two look-ups of the arrays $\gamma$ and $\alpha$.

**Fourth step: final sorting** In the fourth step, positions with identical $l$-mer sequence, grouped together in adjacent positions in $\alpha$ by the previous step, can be further sorted in parallel by longer sequences of length $k > l$ using two-letter bases from positions $l + 1$ to $k$. This sorting step allows fingerprints of any length $k'$ in reads to be retrieved by performing binary search in the subsequent $k' - l$ positions after direct look-up. We use a radix sort algorithm using each of the $k - l$ subsequent bits. The value of $k$ therefore denotes the maximum value for which exact matches under two-letter encoding are queried during the mapping step.

Finally, the vectors $\gamma$ and $\alpha$ are saved to disk, along with a compressed version of the original genome using four bits per base. The count vector $\gamma$ has size $2^l$, and the index vector $\alpha$ has an expected size of $2(M - k)/(W + 1)$. The original genome is stored in a compressed format, where each base (A, C, G, T and N) is saved using four bits per base (Table 1). Since $\gamma$ and $\alpha$ store unsigned 32-bit or (4-byte) integers, the resulting size of the abismal index, in bytes, is expected to be $4(2^l + 2(M - k)/(W + 1)) + M/2$, and is at most $4(2^l + M - k) + M/2$. The factor $2/(W + 1)$, which multiplies the genome size, is a well-established expected value of indexed bases in minimizer theory [4]. For instance, indexing the human genome ($M \approx 3 \times 10^9$) with $l = 24$, $k = 128$ and $W = 13$ requires approximately 3.5 GB. We can verify that, for the tested reference genomes, the resulting size attains the expected size from the formula, that is, only about one in every $(W + 1)/2$ base is indexed. Creating an index genome file can be done using the `abismalidx` program, but users can also provide an input reference genome as a FASTA file directly using the `abismal` program, which is used to map reads to an indexed genome. In this mode, abismal will first index the genome prior to starting to map reads and will not save an intermediate file to disk. It takes about five minutes to index the human genome on a 16-core computer, and about eight minutes using only one thread.

**Estimation of hit cut-off from fingerprint statistics** In the second indexing step (determining fingerprint frequencies), abismal estimates a genome-specific cut-off $c$ for the maximum number of acceptable hits from each seed sampled by reads. To do this, a copy of the vector $\gamma'$ is created and sorted, and a fixed quantile $\beta$ is used to select the cut-off. Specifically, the element in position $\lfloor 2^l(1 - \beta) \rfloor$ of the sorted copy of $\gamma'$ is used as the cut-off estimate. We use $\beta = 10^{-5}$, which means 99.999% of the $l$-mers are used and 0.001% are skipped if selected from reads. This procedure skips certain non-informative $l$-mers such as poly-purines and poly-pyrimidines, which are ubiquitous in many animal genomes and often not helpful for filtration. This value is also saved in the index to be used as default, but abismal can also be run in a full sensitivity mode that sets $c = \infty$ (or, alternatively, $\beta = 0$) regardless of genome fingerprint statistics. Note that no genome positions are excluded from the index based on this number, so this value can be adjusted as necessary if repetitive $k$-mers must be used to map reads.

#### 1.2 Single-end mapping

Reads are mapped in two steps (1) the Hamming distance step collects likely candidates based solely on mismatches, and is further subdivided into a sensitive and a specific step (2) the local alignment step performs banded Smith-Waterman alignments between the read and the candidates collected in the first step. In what follows we will use $k$ to refer to the $k$-mer size of contiguous subsequences used for filtration in the first step.

##### 1.2.1 First step: Hamming distance candidate collection

The Hamming distance step retrieves genome candidates using the vectors $\gamma$ and $\alpha$ calculated during indexing. We assume an input read $r$ is given. A candidate retrieval of a substring of length $k$ starting at position $i$, for $0 \leq i \leq |r| - l$, retrieves all exact matches in the genome of the two-letter encoding of the contiguous substrings selected from $r$. The Hamming distance step is further subdivided into two more steps (1) a specific steps that seeks exact matches, and a (2) sensitive step that performs more comparisons accounting for possible mismatches and indels.

**Specific Hamming distance step** In the specific step, we set $k = 128$ and use the two-letter substrings starting at position $i$ and ending at position $\min(i + k - 1, n - 1)$ for $i \in \{0, \ldots, W - 1\}$. This procedure guarantees that every exact match is collected if one exists in the reference, as the maximum number of non-indexed positions between two indexed positions is $W - 1$. Positions are retrieved through direct look-ups of the $\gamma$ and $\alpha$ vectors for the first $l = 24$ bases, then binary searching the remaining $k - l = 104$ bases. This procedure collects all two-letter exact matches between read and reference.

**Sensitive Hamming distance step** In the sensitive step, we apply the same minimizer selection procedure in read $r$ as we did in the indexing step of the reference genome, where, for $0 \leq r < |r| - W - l$, we select one $l$-mer as the minimizer of each window of size $W$. This procedure selects $k$-mers at most $W = 13$ bases apart, and in average $(W + 1)/2 = 7$ bases apart. Vectors $\gamma$ and $\alpha$ are used to retrieved the indexed positions from the genome. For $l$-mer $v$, the exact two-letter matches indexed in the genome are located in positions starting at $\alpha(\gamma_v)$ and ending at $\alpha(\gamma_{v+1} - 1)$. If any $k$-mer retrieves more than $c$ candidates, the $k$-mer is skipped and the read is aligned based on other $k$-mers.

In both sensitive and specific steps, the Hamming distance between the read and the collected hits is counted. When counting mismatches, Ts in T-rich reads are allowed to match to both Cs and Ts, and As in A-rich reads are allowed to match both As and Gs. Every other base must match the reference exactly. When calculating Hamming distance, abismal requires the number of mismatches to the read to be below a certain value for a candidate location to be considered a valid hit and subsequently align it. We set this value as $Fn$, where $F = 0.4$ by default, that is, each candidate must match at least 40% of the read's bases. This is motivated by the fact that, for typical values of $n$ originating from short Illumina reads, *i.e.*, $50 \leq n \leq 150$, the expected relative number of mismatches of two *i.i.d.* sequences with $\Pr(\text{A}) = 0.3$, $\Pr(\text{G}) = 0.2$ and $\Pr(\text{T}) = 0.5$, conditioned on an identical two-letter match of size $k = 32$, is greater than 0.4 with $> 90\%$ probability. This can verified by simulating a large number of random reads for all possible values of $n$. If a read matches a candidate significantly but contains indels, there is a high probability that the number of matches will be above the 40% threshold, as at least one contiguous substring will contain predominantly matches to the candidate. We keep up to $v = 20$ valid hits for subsequent alignment, but we favor hits with fewer mismatches. To attain this, for each read, we keep a max heap, where triplets $(p, d, s)$ are stored in each element. Here $p$ is the position in the genome, $d$ is the Hamming distance and $s$ is the strand. At the root of the max heap is the candidate with highest Hamming distance, and when a new candidate is compared, if the Hamming distance is below the root's distance ($d$), then the element is added to the heap and the root is discarded if the heap size is above $v$. In single-end mapping, if at least two exact matches are found, then the comparison is stopped as the read is guaranteed to be ambiguous.

**Hamming distance step for paired-end reads** Abismal maps paired-end reads by assuming that the best concordant pair is the most likely mapping location. Two reads can map ambiguously to several locations in the genome, but if only one pair of such locations is concordant in paired-end mapping, the read is said to map uniquely. Conversely, two read pairs can have equal similarity, with reads from individual ends having unique mapping locations. For this reason, in paired-end mapping, we continue to collect hits even when several exact matches are found.

After finding the best approximate matches independently and populating each read end's heap, we test all pairs of heap elements for concordance. The maximum heap size for paired-end mapping is $v' = 200$. Given two reads $r_1$ and $r_2$ whose local alignments to the genome start and end at positions $P_1 = (s_1, e_1)$ and $P_2 = (s_2, e_2)$, respectively, where $s_1 \leq e_1$ and $s_2 \leq e_2$, we say $P_1$ and $P_2$ are concordant if they map to the same chromosome and $d \leq e_2 - s_1 \leq D$, where $d$ and $D$ can be user-defined as minimum and maximum fragment sizes and set by $d = 32$ and $D = 3000$ by default. Note that the fragment length can be smaller than individual read lengths, that is $e_2 - s_1 < \min(e_1 - s_1, e_2 - s_2)$. These "dovetail reads" are infrequent but possible in WGBS datasets, and therefore accepted by abismal. The best pairs are chosen independently of fragment size and solely based on the sum of alignment scores of the two fragments. Concordance tests between pairs of candidates are performed by sorting candidates from both ends by genome position $p$. If the best concordant pair has one unique end and one ambiguous end, only the unique end is reported. If no concordant pairs are found, abismal reports the end which highest alignment score if its edit distance is below the acceptable cut-off. Abismal favors paired-ended maps over single-ended maps even if the ends in the best pair have more mismatches than the any end in the best single-ended maps.

##### 1.2.2 Second step: Local alignments

The $v$ best candidates retrieved in the approximate matching step (in single-ended mapping) or in all concordant pairs (in paired-ended mapping) are aligned to the reference genome if the number of mismatches is less than $Fn$. Local alignments are performed using the banded Smith-Waterman algorithm with a band width of 3, meaning that no reads are allowed to have more than 3 consecutive insertions or deletions. We use a scoring system of +1 for matches and -1 for mismatches and indels.

#### 1.3 Reporting mapped reads

The positions with highest local alignment score are selected. Reads are reported as unique if the candidate that maximizes alignment score (single-end mapping) or the sum of alignment scores (paired-end mapping) is unique and has edit distance below the fraction $f$ of the read length (single-end) or if both ends are simultaneously below a fraction $f$ of their read lengths. We set $f = 0.1$ by default, but this value can be defined by the user if more or less error is acceptable. Reporting locations for ambiguous reads can lead to incorrect methylation estimates in CpGs downstream of read mapping, but may be needed for application-specific global statistics on a dataset. By default abismal does not report ambiguous reads, but users can choose to output a random location using the boolean `-A` parameter, in which case ambiguity is reflected in the SAM flags.

#### 1.4 Mapping all possible combinations of T-rich and A-rich reads

The framework above is used to map both traditional and RPBAT reads in both single-end and paired-end mapping. Mapping traditional single-end reads assumes the input to be T-rich and its reverse-complement (when mapping to the reverse strand of the genome) to be A-rich. Mapping single-end PBAT reads or the second end of paired-end reads assumes the input read is A-rich, and maps its reverse-complement as T-rich. Single-end random PBAT datasets are mapped by trying all possibilities, that is, first mapping the

Table 1: Four-bit encoding of each nucleotide letter

| | A | C | G | T | N |
|---|---|---|---|---|---|
| **reference encoding** | 0001 | 0010 | 0100 | 1000 | 0000 |
| **read encoding (T-rich)** | 0001 | 0010 | 0100 | 1010 | 0000 |
| **read encoding (A-rich)** | 0101 | 0010 | 0100 | 1000 | 0000 |

original read as T-rich (and its reverse-complement A-rich), then mapping the original read as A-rich (and its reverse-complement as T-rich). The information for strand and bisulfite base is kept for all candidates. For every bisulfite sequencing protocol, the strand is reported through SAM flags, and bisulfite conversion is reported through an additional SAM tag which displays `CV:A:T` for reads mapped assuming T-rich and `CV:A:A` for reads mapped assuming A-rich. Mapping paired-end random PBAT datasets is done similarly to single-end, but corresponding reads in read pairs are always assumed to have complementary bisulfite bases, so abismal always reports corresponding reads in pairs with complementary `CV` tag values.

##### 1.5 Implementation choices

Because the goal of abismal is to map bisulfite-converted reads efficiently, many implementation choices were made to accelerate certain critical parts of the read mapping routine. We specifically highlight our choice to encode if a reference base and a read base are a match or mismatch. For both read bases and reference bases, we encode each letter using four bits per letter. The binary encoding of each base is highlighted in Table 1. This encoding has the following property: read base `x` and reference base `y` are a match if and only if `x&y` have exactly one active bit. We use this property to efficiently count mismatches between encoded read and reference sequences. Our encoding of the reference genome groups 16 consecutive letters in 64-bit integers, using 4 bits per letter, and a total of $\lceil M/16 \rceil$ integers. We encode reads similarly, using $\lceil n/16 \rceil$ integers per read. For read integer `w` and genome integer `u`, the number of mismatches between the two sequences is the number of active bits in `w&u`, which can be efficiently counted using built-in registry operations (specifically, we use the `__builtin_popcountll` function in C++).

#### 2 List of abismal constants

Many constant values are used in the abismal algorithm, as described in Section 1. The value for these constants was selected through simulations from the human genome. Each parameter was chosen to maximize sensitivity in the algorithm up to a point where the increase in mapping time and memory does not compensate the increase in sensitivity.

Table 2: List of constants used in the abismal algorithm

| parameter | symbol | value | description and effect of parameter change |
|---|---|---|---|
| window size | $W$ | 13 | larger windows reduce index size and mapping time but may reduce sensitivity given the minimizer property that only guarantees seed to be found when there is an exact match of size $W + l - 1$. If $W = 1$ the entire genome is indexed. Increasing $W$ reduces the index size by an expected factor of $2/(W + 1)$. |
| seed size | $l$ | 24 | larger seeds make the algorithm faster but less sensitive to two-letter mismatches and indels. A read must have at leasts one minimizer that is an exact match to a reference minimizer to be correctly mapped. |
| $l$-mer cut-off | $\beta$ | $10^{-5}$ | A quantile is specified as a cut-off estimate for the maximum number of seed hits when $k$-mers (or $l$-mers) are sampled from reads. Setting $\beta = 0$ does not exclude any seed sampled from reads, and a much larger number of Hamming distance comparisons are performed based on potentially non-informative $k$-mers. Therefore, decreasing $\beta$ increases mapping accuracy at the expense of larger mapping times. |
| hit cut-off | $F$ | 0.4 | A hit is only considered for alignment if the Hamming distance is below $Fn$, where $n$ is the read length. If $F$ is too large, the search space increases. Therefore, the accuracy of the algorithm, increases, but a significantly larger number of alignments is performed to find the best hit, which also increases the total mapping time. |
| SE heap size | $v$ | 20 | The $v$ candidates with lowest Hamming distance are kept in a max heap and subsequently aligned using a banded Smith-Waterman algorithm. Increasing $v$ increases both accuracy and run time. |
| PE heap size | $v'$ | 200 | The $v'$ candidates with lowest Hamming distance for each end of a read pair are kept in a max heap. Each of the $v' \times v'$ pairs of elements in both heaps are then tested for concordance, that is, if they fall within minimum and a maximum insert size. Increasing this value makes it more likely that a read pair will be found but also increases the average number of alignments performed and the time to update the heap at each new valid hit. |
| band width | $b$ | 3 | higher bandwidth will increase sensitivity as a larger alignment is performed which increases the likelihood of passing an alignment score cut-off. For a read of length $n$, each requires filling a table with $(2b + 1)n$ cells, so larger alignments will also increase mapping time. |

### 3 Benchmarking

Tests were performed on a set of Intel Xeon CPU processors with 2.60 GHz clock speed and 16 cores. All machines had a CentOS Linux operating system installed. To perform time comparisons, mapping commands were executed by instructing computers to use one node and 16 threads per node, and FASTQ files were read using local disk to avoid variability in IO. We used a snakemake workflow [2] to uniformly process every tested FASTQ file across all used programs. The same parameters were used to map simulated reads and public datasets.

To the extent allowed by each program, parameters were standardized so that the algorithms behave similarly. The number of mismatches was set as 10% of the read length for single-end samples and 10% of the read length of each end for paired-end samples. The minimum and maximum insert sizes for concordant pairs were set to 32 and 3000, respectively, using mapper-specific parameters. All mappers we instructed to use 16 threads for mapping, output only unique reads and bypass trimming steps prior to mapping. When any of these properties could not be set by parameters, custom scripts were written to further adjust the mapper output for downstream analyses

**Read simulation** Sherman ( https://github.com/FelixKrueger/Sherman ) was used to simulate bisulfite-converted reads. We ran the program with the `-l` flag used to specify read lengths from 50 to 150 in steps of 10. We also set the `-pe` flag, and the `-n` flag was set to 1000000 to simulate one million paired-end reads for each of the six error rates simulated. We set the `-I` flag to 32 as the minimum insert size and the `-X` flag to 500. The `-CH` flag was set to 99 to simulate 99% conversion rate in non-CpGs, and the `-CG` flag set to 30 to simulate 70% methylation in CpGs. The `-e` flag was used to set error rates to 0, 1, 2, 3, 4 and 5.

**Public data downloading and preprocessing** Reads were downloaded using the `fastq-dump` program from the SRA toolkit. Adapter trimming was performed using the `trim_galore` program, which was instructed to trim only Illumina adapters and keep all other bases using the `-q` flag to 0. We set the minimum read length post-trimming as the program default, which is 20 bases. Reference genomes for human (hg38), mouse (mm10), chicken (galgal6), zebrafish (danre11), chimpanzee (pantro6) were downloaded from the UCSC genome browser database [1]. For human and mouse, only primary assembly chromosomes were used. The arabidopsis genome (tair10) was downloaded from the arabidopsis information resource [3].

**Parameters for abismal** Abismal version 1.0.0 was run with all optional parameters set to their default values as described in section 2, with the exception of the `-t` parameter, which was set to 16 to map reads in parallel using 16 threads. We ran abismal by passing the FASTA reference genome file directly through the `-g` flag, thus requiring the genome to be indexed prior to starting to map reads.

**Parameters for Bismark** Bismark version 0.22.3 was run with the `--parallel` flag set to 8, which split files into 8 parts and ran eight parallel bowtie processes, each running in two cores. The `--icpc` flag was used to bypass printing annotations in FASTQ name. The `-I` and `-M` flags were used to define paired-end insert sizes, and the `--non-directional` flag was used for RPBAT datasets. A custom script was written to standardize the edit distance reported by Bismark in the NM flag. Bismark considers converted bisulfite bases as mismatches relative to the reference genome, so the edit distance was recalculated by subtracting methylation-specific mismatches reported in the XM tag, which displays the bases considered to be unmethylated in and out of the CpG context. Alignment scoring parameters were set on bowtie2 directly. The match and mismatch scores, set by parameters `--ma` and `--mp`, respectively, were both set to 2. Gap open and extension for both insertion and deletions were set to 2 by passing the value 2,2 to parameters `--rdg` and `--rfg`. This essentially sets the alignment scoring scheme to +2 for matches and -2 for mismatches and indels, which is similar to abismal's scoring but scaled by a factor of 2. The minimum alignment score for valid reads was set to 1.6 times the alignment length by setting the `--score-min` flag to L,0,1.6. This is equivalent to accepting reads whose alignment score under the 1,-1,-1 scheme is 90% of the aligned read length.

**Parameters for BSMAP** BSMAP version 2.90 was run with the `-p` flag set to 16 to instruct the program to run using 16 threads. The `-r` parameter set to 0, which disables reporting reads that map ambiguously. The `-v` flag was set to 0.1, instructing 10% mismatches to be allowed per read, and the `-f` flag was set to 0, avoiding low-quality bases to be trimmed. Gapped alignment was enabled by setting the `-g` flag to 3. Random PBAT data sets were mapped with the `-n` parameter set to 1. The `-m` and `-x` flags were used to select minimum and maximum insert sizes, respectively.

**Parameters for BWA-meth** BWA-meth version 0.2.2 was run with the `--threads` parameter set to 16. We modified the function `bwa_mem` in the python script on file `bwamem.py` to add alignment score parameters. We set parameters `-A`, `-B`, and `-E` to 1 to set match, mismatch, and gap extension penalties to 1. The `-O` flag was used to set gap open penalty to 0. We set the `-L` parameter to 0 to not penalize soft-clipping. A custom script was written to parse the BWA-MEM outputs to only keep reads that were successfully mapped and whose edit distance, reported in the `NM` tag, was below 10% of the read length. Mapping statistics were obtained through the `samtools stats` program before filtering reads.

**Parameters for HISAT-3N** HISAT-3N version 2.2.1-3n was run with the `--no-spliced-alignment` option to by pass split read alignment. the `--base-change` parameter was set to `C,T`. Insert size parameters `-I` and `-X` were set to 32 and 3000. Parameters `--mp`, `--sp`, `--np`, `--rdg` and `--rfg` were all set to 1 to make alignment scores similar to that of all other mappers. Gap open penalties were also set to 0 using the `--rdg` and `--rfg` flags. The `-score-min` parameter was set to `L,0,-0.1` to set a 10% edit distance cut-off based on the alignment scoring scheme.

**Parameters for WALT** WALT version 1.0 was run with the `-t` (number of threads) parameter set to 16. and the `-P` flag used for RPBAT datasets. The number of mismatches was set as 10% of the read length of each dataset using the `-m` flag. The `-L` flag was used to set the maximum insert size. A custom program was written to convert the MR output from WALT to SAM for further downstream analysis.

**Concordant pair measurement** The percentage of total and unique reads mapped by each program was obtained by parsing text reports of the number of total and uniquely mapped reads generated by each mapper.

**Error rate measurement** Error rates were calculated using the `samtools stats` program. which computes the ratio between the edit distance reported in the NM tag and the total number of mapped bases, calculated by the CIGAR strings.

**Methylome analysis** Bisulfite conversion, CpG coverage and CpG depth were quantified using the methpipe suite [5] version 5.0.0. Paired-end SAM files were converted to single-end by merging mates using the `format_reads` program. Duplicate reads, defined as reads that map to identical location and strand in the genome with equal lengths, were removed to avoid biases in methylation quantification as a result of PCR over-amplification. Bisulfite conversion rate was calculated using the `bsrate` program. The number of converted and unconverted cytosines in reads covering each genome cytosine was quantified using the `methcounts` program. Coverage, depth, and methylation levels for for cytosines and CpGs were quantified using the `levels` program.

The testing pipeline, which displays the commands used to run each program and the memory and number of threads allocated to each, is available at github.com/guilhermesena1/abismal_benchmark . The repository also contains the output files generated at each command executed, the scripts used to parse output files from each program and generate figures and tables displayed in this manuscript, as well as the custom software tools used to format mapper output files when necessary.
